# Priming mycobacterial ESX-secreted protein B to form a channel-like structure

**DOI:** 10.1101/2021.01.02.425093

**Authors:** Abril Gijsbers, Vanesa Vinciauskaite, Axel Siroy, Ye Gao, Giancarlo Tria, Anjusha Mathew, Nuria Sánchez-Puig, Carmen López-Iglesias, Peter J. Peters, Raimond B. G. Ravelli

**Author notes:** European Institute of Chemistry and Biology (IECB), Pessac, France. Dipartimento di Chimica “Ugo Schiff”, Università degli Studi di Firenze, Via della Lastruccia, 3-13 I-50019 Sesto Fiorentino, Italia (Giancarlo Tria). Corresponding author (RBGR), (PJP).

## Abstract

ESX-1 is a major virulence factor of *Mycobacterium tuberculosis*, a secretion machinery directly involved in the survival of the microorganism from the immune system defence. It disrupts the phagosome membrane of the host cell through a contact-dependent mechanism. Recently, the structure of the inner-membrane core complex of the homologous ESX-3 and ESX-5 was resolved; however, the elements involved in the secretion through the outer membrane or those acting on the host cell membrane are unknown. Protein substrates might form this missing element. Here, we describe the oligomerisation process of the ESX-1 substrate EspB, which occurs upon cleavage of its C-terminal region and is favoured by an acidic environment. Cryo-electron microscopy data are presented which show that EspB from different mycobacterial species have a conserved quaternary structure, except for the non-pathogenic species *M. smegmatis*. EspB assembles into a channel with dimensions and characteristics suitable for the transit of ESX-1 substrates, as shown by the presence of another EspB trapped within. Our results provide insight into the structure and assembly of EspB, and suggests a possible function as a structural element of ESX-1.

## Introduction

Tuberculosis (TB) is an infectious disease caused by the bacillus *Mycobacterium tuberculosis*. It is estimated that one-quarter of the world’s population is currently infected with latent bacteria. In 2018,10 million people developed the disease from which 0.5 million were caused by multidrugresistant strains. Even though TB is curable, 1.5 million people succumb to it every year (World Health Organization, 2019). The current treatment is long and with serious side effects, often driving the patient to terminate the therapy before its conclusion (Schaberg *et al.*, 1996). This has contributed to an increase in the number of patients suffering from multidrug- and extensively drugresistant TB. While treatment is available for some of these resistant strains, the regimen is usually longer, more expensive and sometimes more toxic. For this reason, research on mycobacterial pathogenesis is vital to find a proper target in order to develop more effective therapeutics and vaccines.

The high incidence of TB relates to the ability of *M. tuberculosis* to evade the host immune system (Ferluga *et al.*, 2020). This ability is related to multiple factors, one of which is a complex cell envelope with low permeability that plays a crucial role in drug resistance and in survival under harsh conditions (Brennan & Nikaido, 1995). Likewise, pathogenic mycobacteria secrete virulence factors that manipulate the environment and compromise the host immune response. Mycobacteria have up to five specialised secretion machineries that carry out this process, named ESX-1 to −5 (together known as the type VII secretion system or T7SS). The core components of the inner-membrane part of T7SS have been identified (Pym *et al.*, 2003). Nevertheless, it remains unknown whether the translocation of substrates through the inner and outer membrane is functionally coupled or not (one-or two-step, respectively), and if it deploys a specific outer-membrane complex to do so (Bunduc *et al.*, 2020a). Proteins from the PE/PPE family, characterised by Pro-Glu and Pro-Pro-Glu motifs and secreted by T7SS, are often associated with the outer most layer of the mycobacterial cell envelope, and have been suggested to play a role in the membrane channel formation (Burggraaf *et al.*, 2019; Cascioferro *et al.*, 2007; Wang *et al.*, 2020). Recently, the intake of nutrients by *M. tuberculosis* was shown to be dependent on PE/PPE proteins, suggesting that these form small molecule-selective porins that allow the bacterium to take up nutrients over an otherwise impermeable barrier (Wang *et al.*, 2020).

ESX-1 to −5 are paralogue protein complexes with specialised functions and substrates, unable to complement each other (Abdallah *et al.*, 2007; Phan *et al.*, 2017). ESX-1 is an essential player in the virulence of *M. tuberculosis.* It has been implicated in phagosomal escape, cellular inflammation, host cell death, and dissemination of the bacteria to neighbouring cells (Abdallah *et al.*, 2011; Houben *et al.*, 2012a; Simeone *et al.*, 2012; Stanley *et al.*, 2007; van der Wel *et al.*, 2007). Our knowledge about the structure of the machinery as well as the mechanism of secretion and regulation remains limited. ESX-3 is involved in iron homeostasis (Siegrist *et al.*, 2009), and only recently the molecular architecture of its inner-membrane core has been determined (Famelis *et al.*, 2019; Poweleit *et al.*, 2019). The complex consists of a dimer of protomers, made of four proteins: ESX-conserved component (Ecc)-B, C, D(x2), and E. Despite the resolution achieved in both studies, there was no obvious channel through which the proteins substrates can traverse. Rosenberg and collaborators have described that one of the elements of the secretion system (EccC) forms dimers upon substrate binding, which then forms higher-order oligomers (Rosenberg *et al.*, 2015). This is in agreement with observations that ESX-5, which is involved in nutrient uptake (Ates *et al.*, 2015) and host cell death (Abdallah *et al.*, 2011), forms a hexamer (Beckham *et al.*, 2020; Bunduc *et al.*, 2020b; Houben *et al.*, 2012b). A recent structure of the ESX-5 hexamer shows that is it stabilised by a mycosin protease (MycP) positioned in the periplasm on top of EccB_5_ (Bunduc *et al.*, 2020b). ESX-2 and ESX-4 are the least characterised, where ESX-4 is involved in DNA transfer (Gray *et al.*, 2016) and is seen as being the ancestor of the five ESX-systems (Gey Van Pittius *et al.*, 2001).

Located in different positions in the genome, the *esx* loci contain the genes that code for the four Ecc proteins, MycP, a heterodimer of EsxA/B-like proteins, and one or more PE-PPE pairs. With high sequence similarity and conservation between paralogues (Poweleit *et al.*, 2019; van Winden *et al.*, 2016), one could expect the inner-membrane core of the different systems to share a similar architecture. So what makes each one of them unique? Experimental data suggest that the answer lies with the substrates (Lou *et al.*, 2017). The *esx-1* locus encodes for more than ten unique proteins that are known to be secreted (Sani *et al.*, 2010), termed the ESX-1 secretion-associated proteins (Esp) (Bitter *et al.*, 2009). Amongst those is EspC, a protein present in pathogenic organisms that was described to form filamentous structures *in vitro* and to localise on the surface of the bacteria *in vivo* (Lou *et al.*, 2017). Due to the similarities between EspC and the needle protein of the type III secretion system, Lou *et al.* hypothesised that ESX-1 could be an injectosome system with EspC as its needle. This is of particular importance because, compared to the other systems, ESX-1 function has been described to take place through a contact-dependent mechanism (Conrad *et al.*, 2017), which makes the discovery of an outer-membrane complex essential for understanding the system. Other proteins, like EspE that has been localised on the cell wall (Carlsson *et al.*, 2009; Phan *et al.*, 2018; Sani *et al.*, 2010), are of interest as possible elements of the outer-membrane complex. The protein EspB has been the focus of attention due to its ability to oligomerise upon secretion (Korotkova *et al.*, 2015; Solomonson *et al.*, 2015), making it a strong candidate as a structural component of the machinery (Piton *et al.*, 2020).

EspB belongs to the PE/PPE family, but unlike other family members that form heterodimers in mycobacteria, EspB consists of a single poly-peptide chain fusing the PE and PPE domains (Korotkova *et al.*, 2015). EspB is a 48-kDa protein that matures during secretion: Its largely unstructured C-terminal region is cleaved in the periplasm by the protease MycP_1_, leaving a mature 38-kDa isoform (Ohol *et al.*, 2010; Solomonson *et al.*, 2013; Xu *et al.*, 2007). The purpose of this maturation is not yet clear but it was shown that inactivation of MycP_1_, and thus cleavage of EspB, deregulates the secretion of proteins by ESX-1 (Ohol *et al.*, 2010). Chen *et al.* observed specific binding of EspB to phosphatidylserine and phosphatidic acid after cleavage (Chen *et al.*, 2013), suggesting that the C-terminal processing of EspB is important for its functioning, possibly involving lipid binding. The crystal structure of the monomeric N-terminal part of EspB from *M. tuberculosis* and *M. smegmatis* has been determined: It forms a four-helix bundle with high structural homology between species (Korotkova *et al.*, 2015; Solomonson *et al.*, 2015). During the preparation of this work for publication, the structure of an EspB oligomer from *M. tuberculosis* was published by Piton *et al.*, showing features of a pore-like transport protein (Piton *et al.*, 2020).

EspB is the only member of the PE/PPE family described to date to form higher-order oligomers. In this work, we studied the oligomerisation ability and structures of EspB from *M. tuberculosis, M. marinum, M. haemophilum* and *M. smegmatis*. We show that truncation of EspB at the MycP_1_ cleavage site and an acidic environment promote the oligomerisation of EspB from the three pathogenic species but not from non-pathogenic *M. smegmatis*. Oligomerisation is mediated by intermolecular hydrogen bonds and amide bridges between residues highly conserved in the pathogenic species, but absent in *M. smegmatis*. The structures of oligomeric EspB consist of two domains: an N-terminal region that forms a cylinder-like structure with a tunnel large enough to accommodate a folded PE-PPE pair, and a partly hydrophobic C-terminal region that interact with hydrophobic surfaces. The oligomer has similar inner-pore dimensions as was described for the pore within the periplasmic region of ESX-5 (Bunduc *et al.*, 2020b). Visualisation of a trapped EspB monomer within the channel supports the idea that it could transit secreted proteins through its tunnel. Overall, in this work we describe factors that prime the oligomerisation of EspB, and provide insight into its potential role in the ESX-1 machinery.

## Results

### Oligomerisation is favoured by an acidic pH and maturation of EspB

Previously, it has been described that oligomerisation of EspB occurs after secretion (Korotkova *et al.*, 2015). In the infection context, this secretion would lead the protein to the phagosomal lumen of a macrophage, an organelle known to have pH acidification as a functional mechanism. To evaluate the putative role of pH in the oligomerisation process, the mature form of *M. tuberculosis* EspB (residues 2-358) was incubated at different pH values and analysed by size exclusion chromatography (SEC). Results showed that the equilibrium is favoured towards an oligomer form at pH 5.5 compared to pH 8.0 (Fig 1A), as observed by a higher oligomer/monomer ratio at any protein concentration (Fig 1B). Native mass spectrometry experiments confirmed this behaviour and could identify different oligomeric states of EspB, with the heptamer being the most predominant.

**Fig 1.**
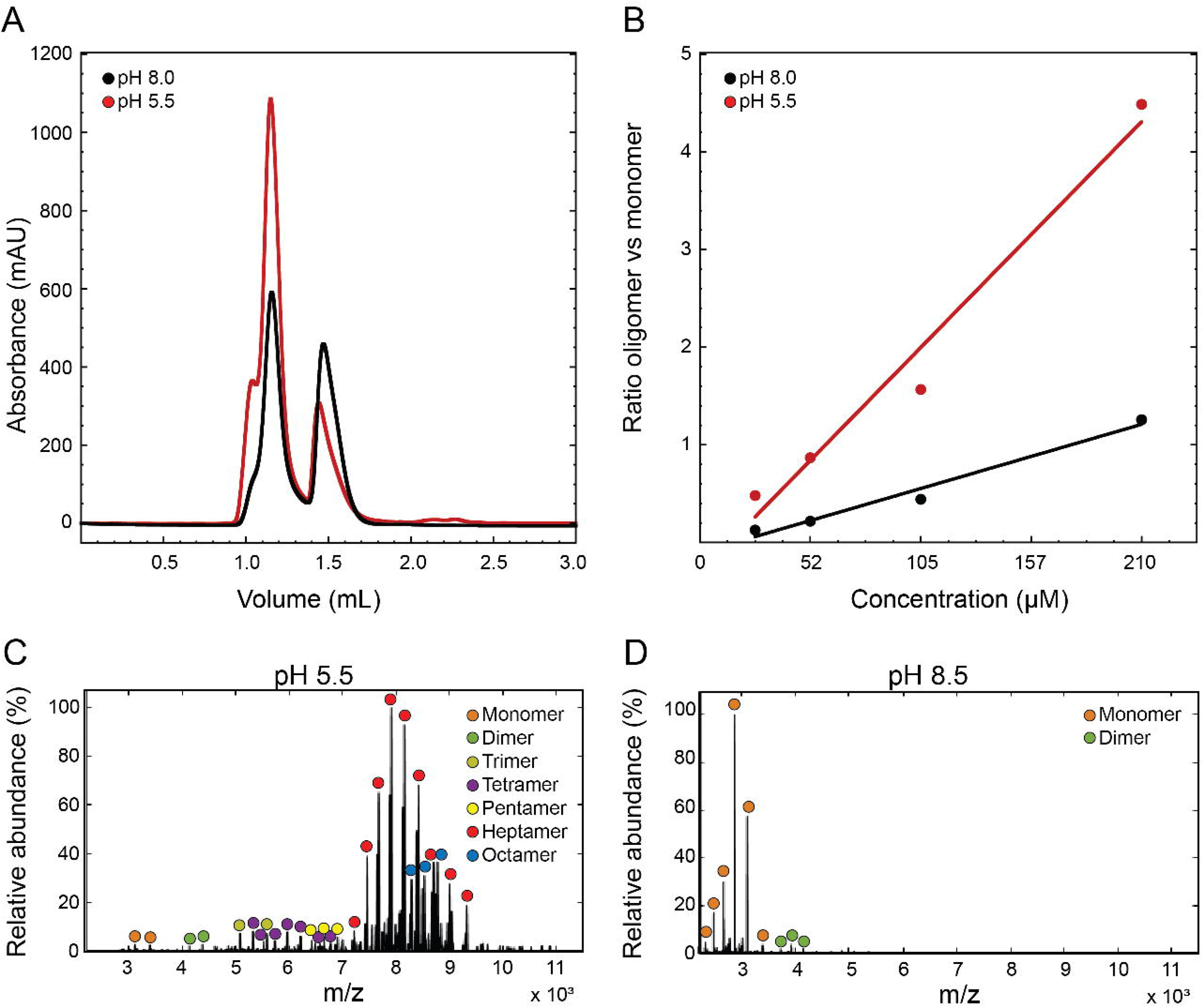
Oligomerisation of EspB is promoted by an acidic environment. (A) Size exclusion chromatography profiles of *M. tuberculosis* EspB_2-358_ at 210 μM in 20 mM acetate buffer (pH 5.5), 150 mM NaCl and 20 mM Tris (pH 8.0), 150 mM NaCl. Void volume corresponds to 0.8 mL elution volume. (B) Oligomer/monomer ratios at different protein concentrations in conditions from panel (A). The absorbance values of the oligomer were taken at 1.14 mL while the monomer values were at 1.48 mL. (C-D) Presence of the different oligomer species from *M. tuberculosis* EspB_2-348_ at pH 5.5 and pH 8.5 obtained by native mass spectrometry.

Intermediate states were observed (dimer to pentamer) and even higher oligomeric states (octamer) but in lower abundance compared to the heptamer (Fig 1C and D).

Because EspB undergoes proteolytic processing of its C-terminus during secretion, we investigated the effect of this cleavage on the quaternary structure of different EspB constructs, varying in their C-terminus lengths, from *M. tuberculosis* at pH 5.5 (Fig 2). With the exception of EspB_7-278_ that did not oligomerise, we observed that oligomerisation was favoured for all other constructs at pH 5.5 (Fig 2B). The full-length EspB_2-460_ (Fig 2B, blue trace) presented the lowest amounts of complex formation compared to the other constructs tested, while the highest amount was observed for the mature isoform, EspB_2-358_ (Fig 2B, orange trace). These results suggest that MycP_1_ cleaves EspB to allow oligomerisation, and that the remaining residues of the unstructured C-terminal region are needed, possibly, to stabilise the complex.

**Fig 2.**
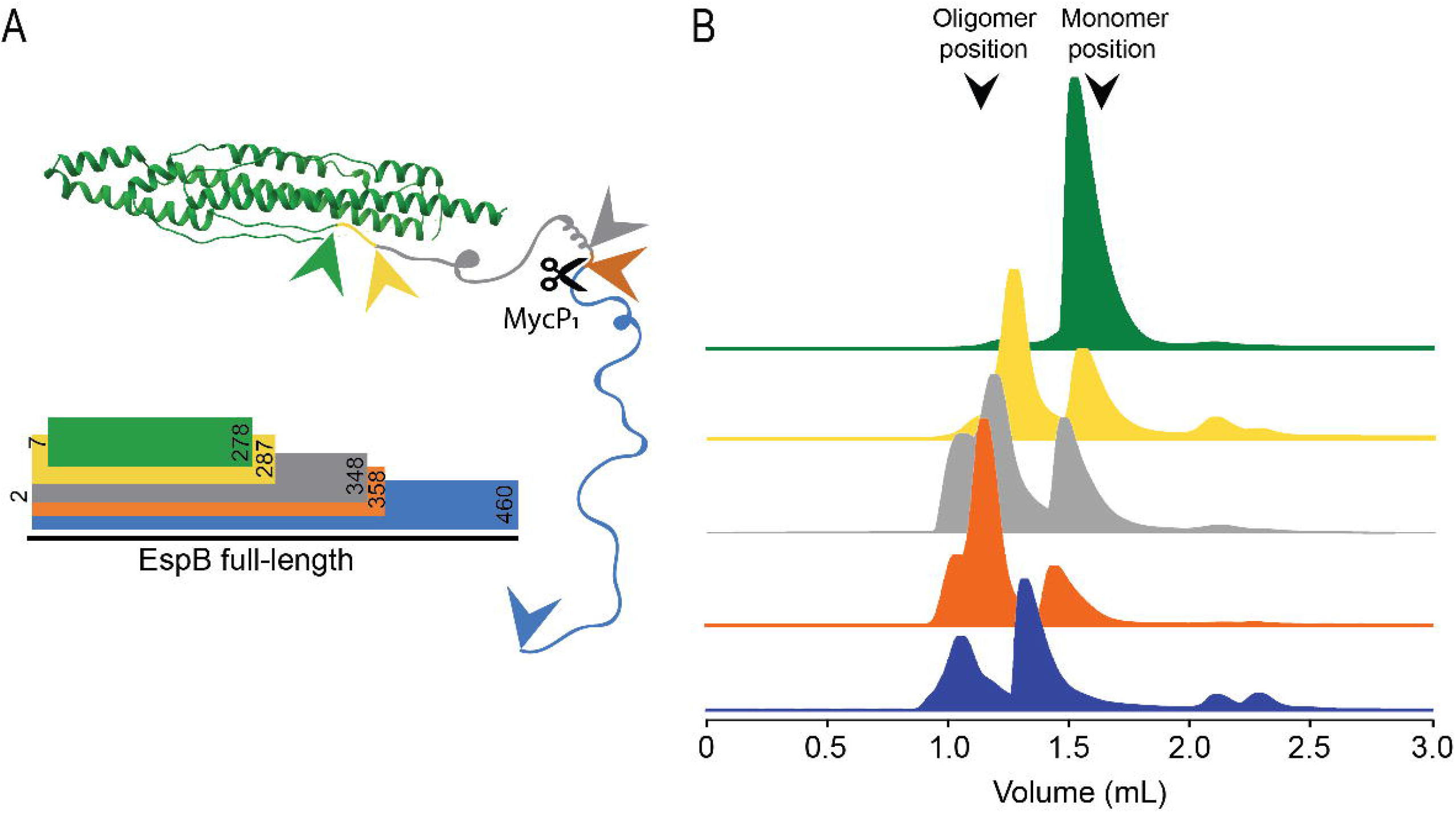
Impact of EspB C-terminus processing on oligomerisation. (A) Scheme of the different constructs used in this work, where EspB_2-460_ is in blue, EspB_2-358_ in orange (MycP_1_ cleavage site), EspB_2-348_ in grey, EspB_2-287_ in yellow and EspB_7-278_ in green. Structural model from PDB ID 4XXX, while the C-terminal region is a representation of an unfolded protein. Arrows represent the end of each construct. (B) Size exclusion chromatograms of each construct corresponding to the colours in panel (A), resulting from 50 μL sample injection at 220 μM eluted in 20 mM acetate buffer (pH 5.5), 150 mM NaCl. Void volume corresponds to 0.8 mL elution volume.

### EspB from *M. smegmatis* is unable to oligomerise

To determine whether the oligomerisation ability is conserved across species, we performed cryo-EM analysis on different orthologues of the mature EspB. Proteins from the pathogenic species

*M. tuberculosis, M. marinum* and *M. haemophilum* were able to oligomerise into ring-like structures while the non-pathogenic *M. smegmatis* did not, as seen by the lack of visible particles (Fig 3A); the structured region of an EspB monomer (30 kDa) has a signal-to-noise ratio too low to be visualised within these micrographs (Henderson, 1995; Zhang *et al.*, 2020). Interestingly, comparison of the tertiary structure from the pathogenic species studied here with the published crystallographic model of EspB from *M. smegmatis* did not show substantial differences (RMSD Cα’s 0.98 - 1.13 Å), apart from an extended α-helix 2 (Fig 3B), absent in our oligomeric structures. To determine whether the differences in oligomerisation ability between *M. smegmatis* and the pathogenic species were due to their primary structure variances, we performed sequence alignment of multiple EspB orthologues. The species that presented oligomerisation showed high sequence identity whereas *M. smegmatis* has the lowest of all (Fig 3C and Fig EV1).

**Fig 3.**
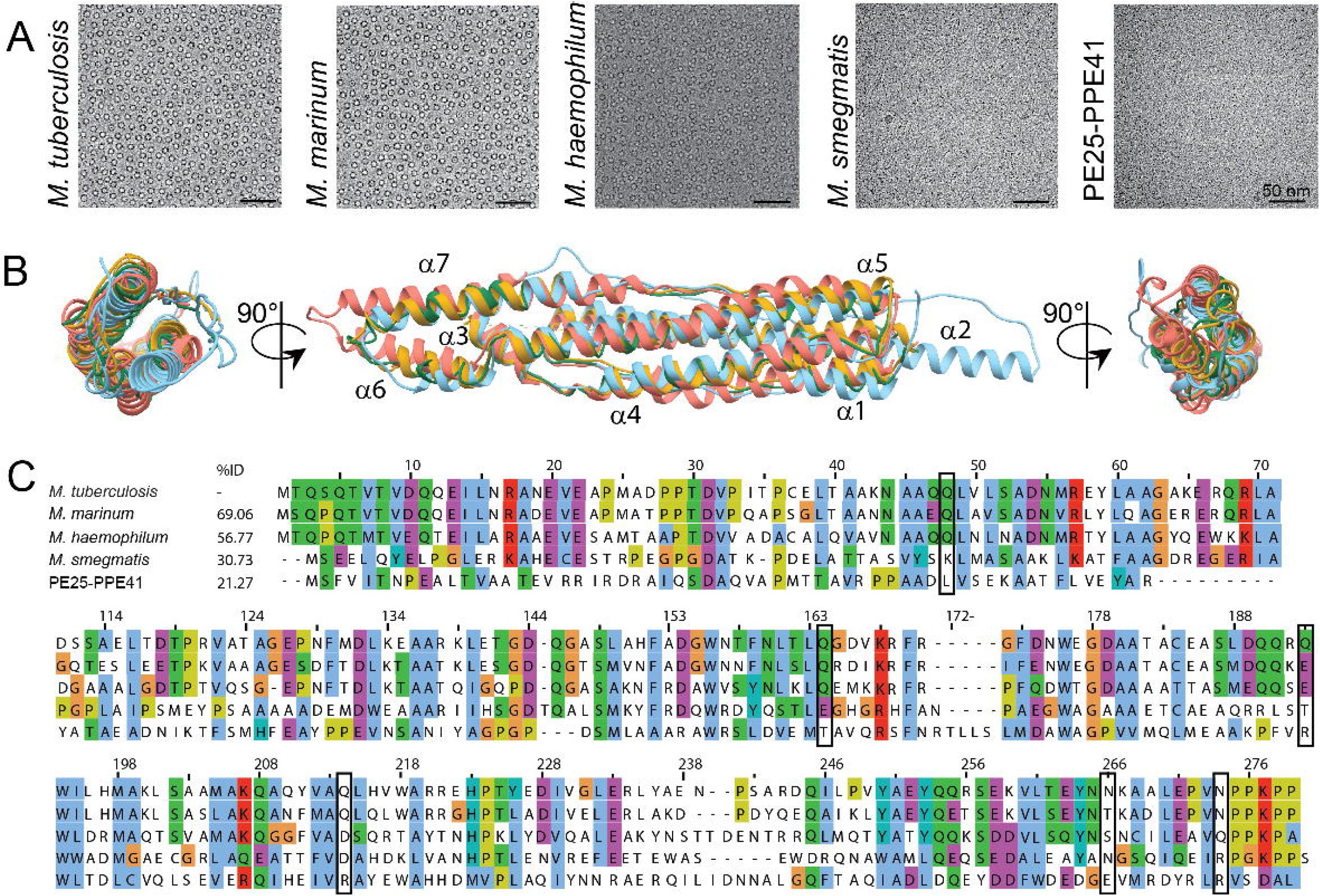
Oligomerisation differences between EspB orthologues despite sharing similar tertiary structure. (A) Evaluation of the oligomerisation of EspB orthologues and PE25-PPE41 by cryo-electron microscopy. Scale bars represent 50 nm. (B) Different views of structural alignment of EspB_Mtb_ (yellow - this work), EspB_Mmar_ (green - this work), EspB_Msmeg_ (light blue - PDB ID 4WJ1), and PE25-PPE41 (orange - PDB ID 4W4K). (C) Multi-alignment of amino acid sequences of different species from the *Mycobacterium* genus, as well as the protein pair PE25-PPE41. Numbering and sequence identity is based on the sequence of *M. tuberculosis.* Rectangles denote residues involved in the oligomerisation of EspB. Alignment was generated using ClustalW server, and figure was created using software Jalview (Waterhouse *et al.*, 2009). The colour scheme of ClustalX is used (Larkin *et al.*, 2007).

Because EspB belongs to the PE/PPE family, we included in the analysis a PE-PPE pair with a structure already published (Ekiert & Cox, 2014). PE25-PPE41 did not oligomerise (Fig 3), despite sharing a similar tertiary structure (RMSD Cα’s 1.134 Å). With a low identity percentage (21.3%), it confirms the importance of specific amino acids sequence for the conservation of the quaternary structure.

### High-resolution cryo-EM structures of EspB oligomers

Next, we aimed to solve the high-resolution structure of EspB oligomers by cryo-EM. Initial experiments were performed with EspB_2-460_ and EspB_2-348_ from *M. tuberculosis*, which displayed a very strong preferential orientation where only “top views” could be seen (Fig 4A). Cryo-electron tomography revealed these molecules to be attached to the air-water interface (Fig EV2). Different oligomers were found: hexamers, heptamers, rings with an extra density in the middle, and octamers, with the heptameric ensemble being the predominant one (Fig 4A), in agreement with the results obtained in solution (Fig 1C). Preliminary 3D reconstructions could be obtained from data that were collected at one or more tilt angles (Fig 4B). Data processing resulted in 3-4 Å resolution maps from which the first heptamer models were built. Removal of C-terminal ending up at residue 287 led to a different distribution of particles on cryo-EM grids, now with random orientations (Fig 4C), implying that this region interacts with the air-water interface on the EM grid (Noble *et al.*, 2018) (Fig EV2).

**Fig 4.**
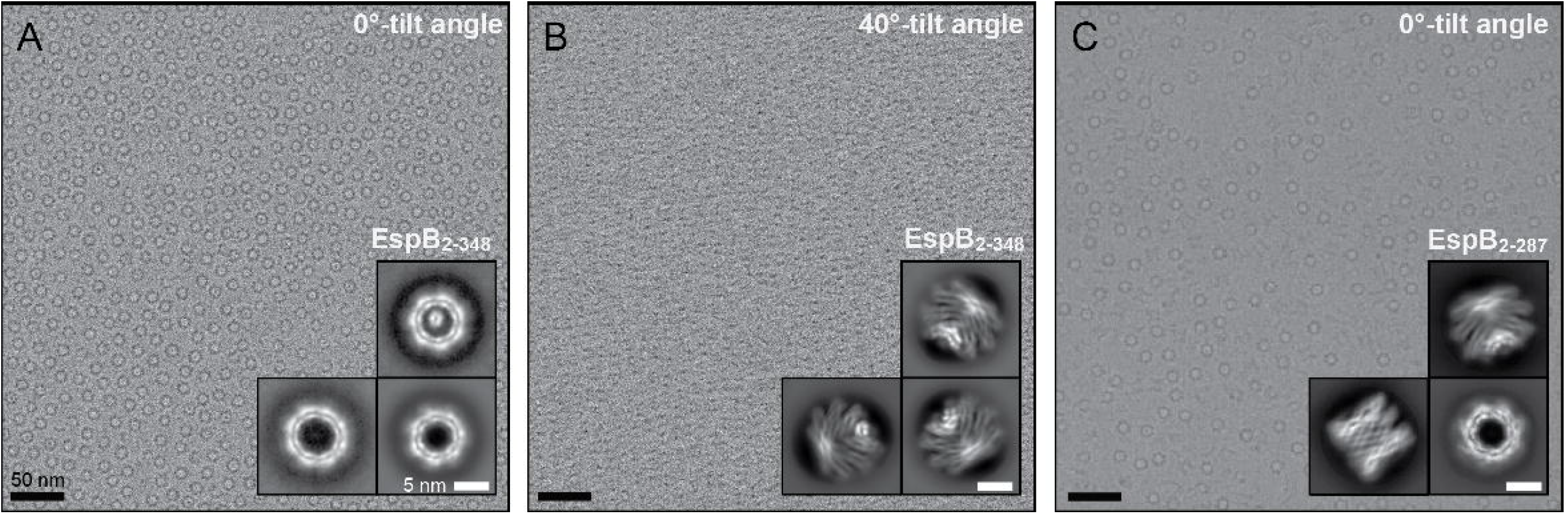
Loss of the EspB preferential orientation by removal of its C-terminal residues. (A-B) Representative micrograph of EspB_2-348_ with preferential orientation at 0°-tilt angle or 40°-tilt angle. (C) Representative micrograph of EspB_2-287_ with random orientation taken at 0°-tilt angle. Insets correspond to the respective 2D classes. Scale bars in A-C represent 50 nm; scale bars in insets represent 5 nm.

Experiments were repeated for constructs EspB_2-287_ from *M. tuberculosis* and the equivalent construct from *M. marinum* (at 0°-stage tilt), leading to high-resolution EM maps of 2.3 Å and 2.5 Å average resolution, respectively (Fig 5A and B, and Fig EV3A-C). We observed high structural conservation between the two structures. Both displayed a four-helix bundle, like the EsxA-B complex and PE25-PPE41 complex, with the WxG and YxxxD located on one end of the elongated molecule, referred to the top hereafter, making an H-bond interaction between the nitrogen of W176 with the oxygen of Y81, as was observed in the crystal structure (Fig 5C) (Korotkova *et al.*, 2015). The helical tip is located on the opposite end, referred to as the bottom, for both EspB and PE25-PPE41 (Korotkova *et al.*, 2015; Solomonson *et al.*, 2015). The C-terminal region starts near the top end of the elongated molecule. The overall structure shows seven copies tilted 32° with respect to the symmetry-axes forming a cylinder-like oligomer with a width and a height of 90 Å (Fig 5A and B).

**Fig 5.**
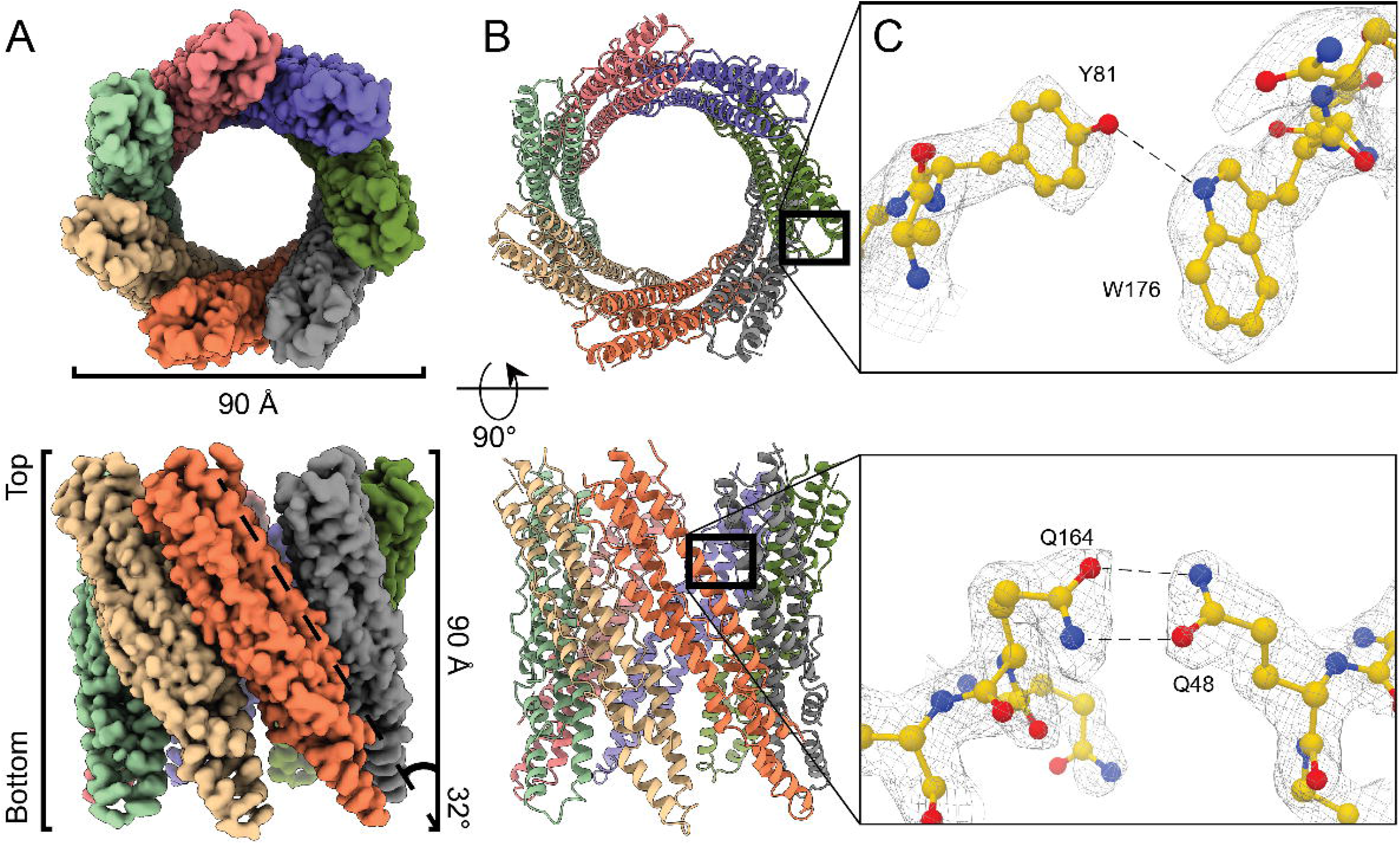
Cryo-EM reconstruction of EspB_2-287_ heptamer complex. (A-B) Density map and structural model made with ChimeraX (Goddard *et al.*, 2018), showing each monomer in different colours. For (A) and (B), the upper panels show the top views and the bottom panels show the side views. (C) Model and densities of intramolecular interaction at W176-Y81 and intermolecular interaction at Q48-Q164. Colours follow the conventional colouring code for chemical elements.

The single particle analysis (SPA) map from *M. tuberculosis* EspB revealed three Q-Q and Q-N interaction pairs between monomers (Fig 5C). Q48 was conserved in all EspB orthologues analysed here with exception of *M. smegmatis* that showed no oligomerisation (Fig 3C and Fig EV1). A Q48A substitution in the *M. tuberculosis* orthologue resulted in the disruption of the oligomer (Fig EV4D, yellow trace) as evidenced by the absence of a high molecular weight peak by SEC. Amide bridges are not commonly seen within the structures present in the Protein Data bank (PDB) (Joosten *et al.*, 2009) but they are stronger interactions than typical hydrogen bonds and less affected by pH changes compared to salt bridges, another strong interaction (Xie *et al.*, 2015). In addition to the Q-Q and Q-N interaction pairs, some hydrophobic interfacing residues were identified, including L51 and L161. Histidines, glutamates and aspartates were not found to interact directly with the neighbouring monomer. It seems that these residues mainly play a role in the pH-dependent overall charge distribution of the monomers (Fig 6A-B).

**Fig 6.**
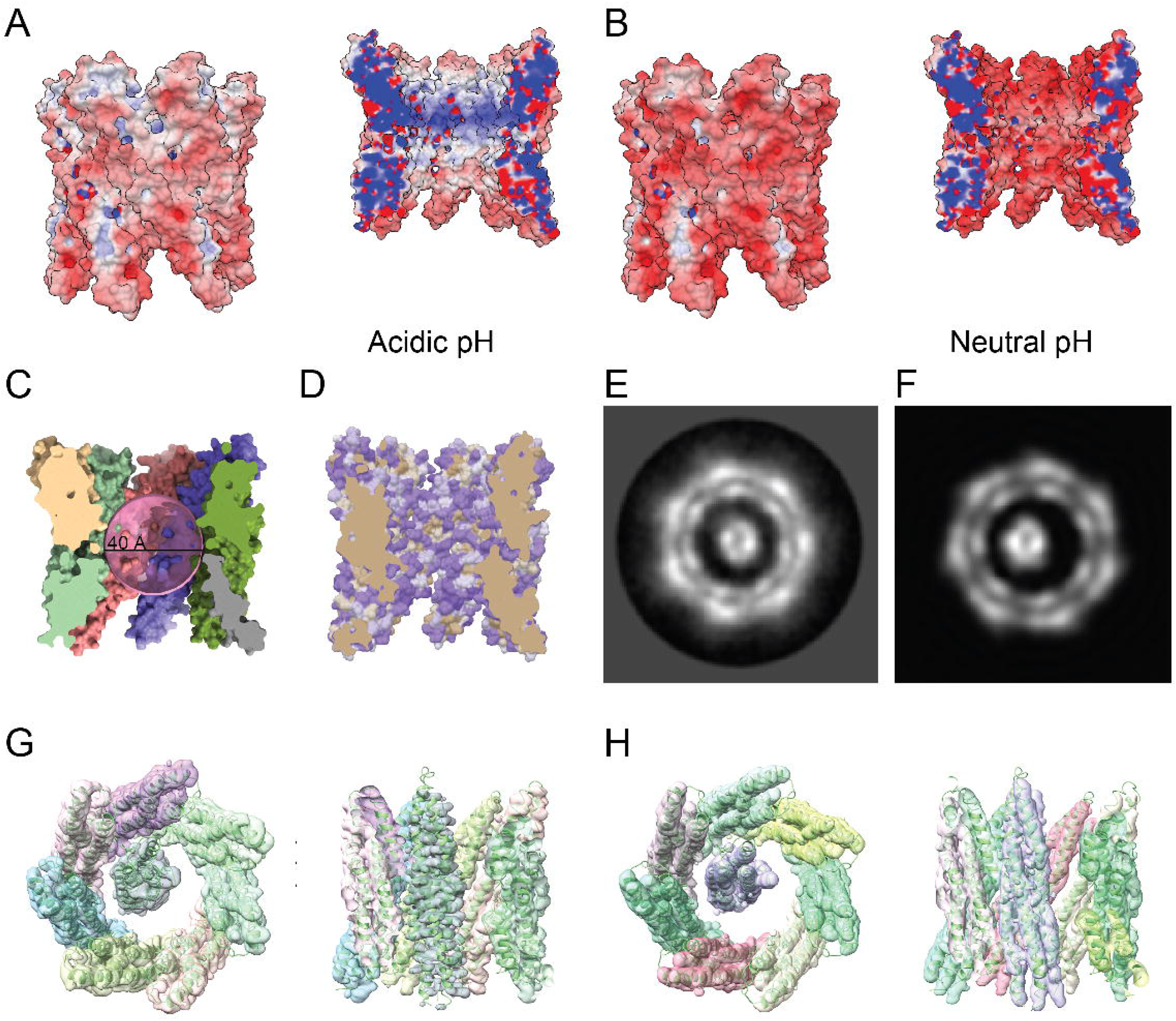
Characterisation of the EspB oligomer. (A) Electrostatic potential of EspB oligomer at acidic and (B) neural pH. The protonation state was assigned by PROPKA (Olsson *et al.*, 2011) and electrostatic calculations were generated by APBS (Baker *et al.*, 2001) and PDBPQR (Dolinsky *et al.*, 2004). (C) The smallest inner diameter of the EspB oligomer is 40 Å, as calculated by HOLE (Smart *et al.*, 1993). (D) Surface representation of amino acid hydrophobicity according to the Kyte-Doolittle scale (polar residues - purple, non-polar residues - gold). (E) High-resolution 2D class of EspB heptamer with extra density in the middle. (F) 2D projection of the 3D map obtained for the 7+1 EspB oligomer. (G) C1 3D map of 7+1 EspB oligomer with local symmetry applied to the heptamer ring. (H) C1 3D map of the 7+1 EspB oligomer with 8-fold local symmetry applied and models fitted to the map.

Our high-resolution EspB oligomer maps did not reveal a continuous density for the PE-PPE linker. The proposed location of the linker within the crystal structure (Korotkova *et al.*, 2015) overlaps with the oligomerisation interface, and would need to adopt a different position upon oligomerisation. Particle subtraction followed by focused classification showed partial densities for the linker at the periphery of the structure. We locked the PE-PPE linker in its crystal-structure position by making a double mutation, N55C (in the core of the monomer) and T119C (in the PE-PPE linker): this would prevent the linker to adopt a different conformation as is needed for the oligomerisation. This double mutant abolished oligomerisation of EspB, suggesting that an intramolecular bond was formed that prevented the linker from moving (Fig EV3D, red trace).

### EspB, a possible transport channel for T7SS proteins

The EspB cylinder-like structure has an internal pore diameter of 40 Å (Fig 6A-B), large enough to accommodate folded proteins such as EsxA/EsxB (diameter 35 Å), PE25-PPE41 (diameter 27 Å) or an EspB monomer itself (diameter 28 Å). Analysis of the degree of hydrophobicity in the structure showed that the internal surface of the oligomer is mainly hydrophilic (Fig 6D), allowing other hydrophilic molecules to pass.

During the cryo-EM data processing, additional densities were consistently found within the EspB heptamer of all the different constructs, including constructs that lack the C-terminal region. Fig 6E shows a high-resolution 2D class of EspB_2-348_ with a well-defined density inside the channel from a subset of the data collected at 0°-stage tilt. This 2D class was found in ~7% of the particles recorded. The 2D classes obtained at 40°-stage tilt could not be unambiguously manually assigned to specific oligomerisation forms. Instead, 3D classification in RELION (Scheres, 2012) was used to identify one class with solely C7 symmetry and one class with an extra density within the heptameric channel. Local symmetry averaging of the heptamer model while processing the overall map in C1 map revealed an extra density spanning the entire channel, in which we could fit an EspB monomer model (Fig 6G-H).

### Integrity of the PE-PPE linker is not essential for the oligomerisation of EspB

To determine if the PE-PPE linker absent in our model was essential for oligomerisation, we performed limited proteolysis analysis on the *M. tuberculosis* constructs. Incubation with trypsin fully digested EspB_7-278_, perhaps due to its lower stability, but resulted in two major fragments for the constructs EspB_2-460_ and EspB_2-348_, as shown by SDS-PAGE (Fig EV4A). N-terminal sequencing and mass spectrometry analysis revealed that the larger fragment corresponded to a section of the protein comprising residues V122 to R343 (corresponding to the PPE domain), while the smaller fragment included the N-terminal end of the protein sequence, with a few residues from the affinity tag, up to residue R121 (PE domain and linker)(Fig EV4B-D). Despite being split within the PE-PPE linker into two fragments, EspB_2-348_ behaved in gel filtration as a single entity with the capacity to form oligomers (Fig EV4E and F) confirming that the integrity of this region is not necessary for the complex to form. It is noteworthy that trypsin did not cut before R343, even though there are cleavage sites in the so-called unfolded C-terminal region rising the question if this region is actually fully unstructured.

### Properties of the EspB C-terminal region

The function of the C-terminal region has puzzled the scientific community for a long time, partly because it is the only substrate known to date of the MycP_1_ protease. Here, we described its processing as an important factor for the oligomerisation of the N-terminal region (1-287), however, the cleavage leaves ~70 residues for no-obvious reason. To gain insight in the properties of the C-terminal region that could hint for its function, and based on the preferential orientation-effect seen in cryo-EM, we performed a hydrophobicity analysis of this region on the different EspB orthologues. Analysis evidenced the presence of hydrophobic patches in the pathogenic species, that are absent in EspB from *M. smegmatis* (Fig EV5A). Some of these patches are present in all the constructs with preferential orientation, leading to speculate that residues 297-324 interact with the air-water interface of the cryo-EM grid (Noble *et al.*, 2018).

To understand whether this effect is related to a structural change or particular characteristic in the C-terminal region of the protein, we expressed a construct corresponding to residues 279-460 and carried out circular dichroism (CD) studies on it. Far UV CD spectra analysis of this region showed a negative band around 198 nm (Fig EV5B), characteristic of random coil structures. This result is in line with the high fraction (54%) of “disorder-promoting” residues within this region (lysine, glutamine, serine, glutamic acid, proline and glycine: amino acids commonly found in intrinsically disordered protein regions). Interestingly, its proline content is 2.5 times higher than that observed for proteins in the PBD (Theillet *et al.*, 2013; Uversky, 2013). Comparative analysis of the CD difference spectra obtained at different pH [Δ□ (pH 5.5 - pH 8.0)] revealed a positive signal close to 220 nm and a negative signal near 200 nm (Fig EV5B inset), showing that this region is able to adopt extended lefthanded helical conformations [poly-L-proline type II or PPII (Rucker & Creamer, 2002)]. CD analysis of the C-terminal region in the presence of different concentrations of 2,2,2-trifluoroethanol (TFE) showed that this region has an intrinsic ability to attain helicity based on the decrease in the ellipticity signal at 222 nm (Fig EV5C) (Luo & Baldwin, 1997). The lack of a single isodichroic point at 200 nm suggests that the conformational changes elicited by TFE do not comply with a two-state model and that most probably the transition is accompanied by an intermediate, *e.g.* the presence of more than one α-helix.

## Discussion

In the present study, we describe different factors that facilitate oligomerisation of EspB: an acidic environment, the truncation of its C-terminal region, a flexible PE-PPE linker and the residues involved in the interaction. Our findings are in agreement with previous observations that EspB oligomerises upon secretion (Korotkova *et al.*, 2015). Based on these results, the C-terminus of the full-length protein could prevent premature oligomerisation in the cytosol of mycobacteria, possibly through steric hindrance. However, this region is also likely to have other functions. Deletion of EspB C-terminus does not affect its own secretion (McLaughlin *et al.*, 2007; Xu *et al.*, 2007) but rather the secretion of EsxA/EsxB, possibly by loss of interaction with the last residues of EspB (Xu *et al.*, 2007). The sequence of the C-terminal end is highly conserved (Fig EV1), which makes it possible that this region interacts with other molecules in the cytoplasm of the bacterium.

This ability of EspB to oligomerise seems to be conserved across mycobacterial species, with the exception of *M. smegmatis.* This microorganism is a fast-growing, non-pathogenic species that uses ESX-1 system for horizontal DNA transfer (Flint *et al.*, 2004). The exact mechanism of this transfer is unknown; however, evidence suggests that ESX-1 is not the DNA conduit but rather secretes proteins that act like pheromones, which in turn induces the expression of *esx-4* genes resulting in mating-pair interactions (Gray *et al.*, 2016). The ESX-1 substrate EsxA was shown to undergo a structural change that allows membrane insertion in *M. tuberculosis* when exposed to an acidic environment, however, this effect does not occur in its *M. smegmatis* orthologue (De Leon *et al.*, 2012; Ma *et al.*, 2015). Taking the aforementioned antecedents and the oligomerisation differences between EspB proteins observed in this work, it is plausible to think that the mechanism of action of the ESX-1 is distinct between these two species.

EspB interacts with the lipids phosphatidylserine and phosphatidic acid (Chen *et al.*, 2013). It was suggested that EspB could transport phosphatidic acid (Piton *et al.*, 2020) but the interior of the complex is mainly hydrophilic, making this scenario less plausible. Despite the presence of lipids in the crystallisation set up, Korotkova *et al.* (2015) could not find lipids within the crystal structure of EspB_7-278_ which lacks the C-terminus. Our results show that the C-terminus of EspB contributes to the protein’s preferred orientation on an EM grid caused by an interaction to the hydrophobic air-water interface (Noble *et al.*, 2018), analogous to what could happen on a lipid membrane. With a PPII helix at the end of the channel followed by hydrophobic patches at the C-terminus, we hypothesise that this secondary structure interacts with the head group of the lipids, as it has been described for other PPII (Franz *et al.*, 2016), allowing the hydrophobic residues to insert into bilayer membranes. Based on the chemical properties of the channel and supported by the evidence of an extra EspB monomer observed within the oligomer, we propose that EspB could be a structural element of ESX-1 allowing other substrates to transit through the channel.

The combined data presented in our work leads us to hypothesise three models of the role of the EspB oligomer. EspB within the cytosol is likely to be monomeric (Korotkova *et al.*, 2015), either free or chaperoned by EspK (McLaughlin *et al.*, 2007). Binding of a chaperone to the helical tip of EspB would place the WxG and YxxxD bipartite secretion signal exposed on the top of EspB, ready to interact with the T7SS machinery. Upon exiting ESX-1 inner-membrane pore, the pre-protein EspB will be cleaved within the periplasm by MycP_1_. Analogous to ESX-5 (Bunduc *et al.*, 2020b), we expect MycP_1_ to cap the central periplasmic dome-like chamber formed by EccB_1_, and to have its proteolytic site faced towards its central pore. The cleavage of the C-terminal region at A358 (Solomonson *et al.*, 2013) will remove the most hydrophilic part of the C-terminus leaving a hydrophobic tail (Fig EV6A).

From here, we propose three possible pathways for the oligomerisation of EspB. In one scenario, after processing of the C-terminus, EspB binds the outer membrane of mycobacteria increasing its critical concentration to form an oligomer (Fig 7 model 1). As suggested above, it can be assumed that EspB monomers would transit through the inner-membrane pore with the top first where the C-terminus as well as the WxG and YxxxD motifs are located. In this position, the monomers would already be properly oriented to form an oligomer on the outer membrane inner leaflet just like EspB_2-358_ attaches to the air-water interface of a cryo-EM grid (Fig 4A). The inner pore of the EspB heptamer has similar dimensions compared to that proposed by Beckham *et al.* for the ESX-5 hexameric structure (Beckham *et al.*, 2017), albeit they later published a higher resolution structure with a more constricted pore in a close state (Beckham *et al.*, 2020). The space between the inner and outer-membrane has been reported to be 20-24 nm wide (Dulberger *et al.*, 2020; Sani *et al.*, 2010; Zuber *et al.*, 2008), which could accommodate the 9-nm long EspB heptamer. It was postulated (Piton *et al.*, 2020) that the positively charged interior of the EspB channel could play a role in the transfer of negatively charged substrates such as DNA or phospholipids. However, in analogy to the negative lumen of a bacteriophage tail that is used to transfer DNA (Zinke *et al.*, 2020), we propose that the positively charged interior space of the EspB oligomer would channel substrates of the same charge, as negatively charged substrates would most likely bind and get trapped. Since the heptameric structure presented here lacks any trans-membrane domains and is highly soluble, it is unlikely to be embedded within the outer membrane but could be anchored by its C-terminus forming part of a larger machinery that completes the ESX-1 core complex. EspB is well known to be secreted to the culture medium of mycobacteria (Lodes *et al.*, 2001), thus in this model EspB will help in its own secretion by forming a channel through which additional substrates like EspB itself, could travel.

**Fig 7.**
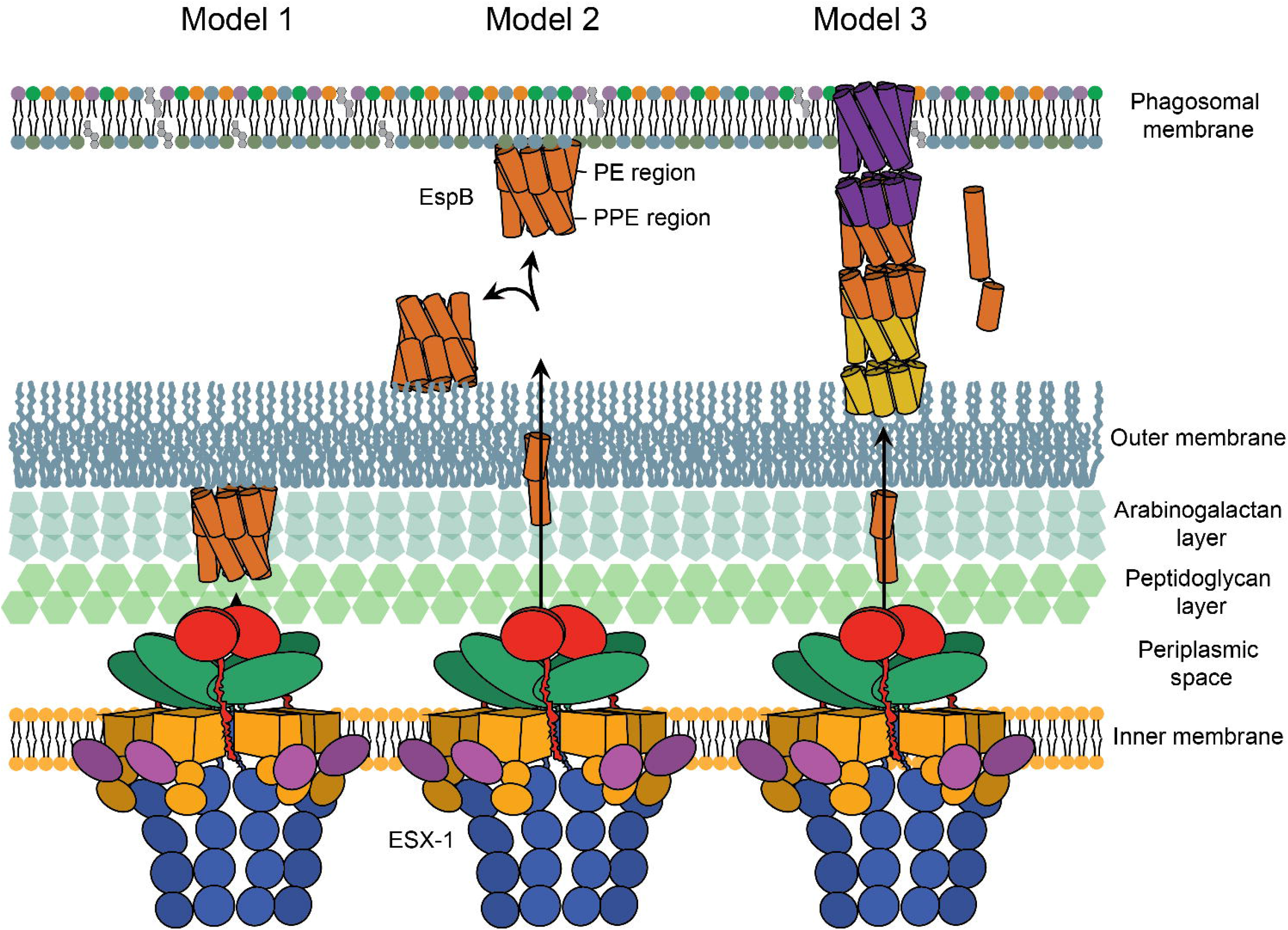
Putative pathways for the oligomerisation of EspB. In Model 1, EspB is cleaved in its C-terminus by the protease MycP_1_ in the periplasm of mycobacteria leaving hydrophobic residues to insert into the outer membrane; an increase in the local concentration on the membrane leads to oligomerisation of EspB. In model 2, secretion of EspB across the double membrane after MyP_1_ cleavage allows the protein to bind to either the phagosomal membrane or the external part of the outer membrane. In model 3, after cleavage in the periplasm and secretion to the exterior of the bacterium, EspB undergoes a conformational change dissociating the PE and PPE domains and exposing hydrophobic residues that would allow the insertion into the membrane; while the PPE gets embedded into the membrane in an oligomeric form, the respective PE is able to interact with the PPE of a second molecule forming a tubular structure. Different colours are used for each heptamer-subunit. Regardless of what oligomerisation pathway EspB follows, oligomerised EspB is hypothesised to form part of the larger machinery that completes the inner-membrane complex of ESX-1.

In a second scenario, after EspB got secreted outside the bacterium, it could interact with either the phagosomal membrane or the external face of the outer membrane (Fig 7 model 2). The aforementioned hypothesis of how EspB is secreted (C-terminus and WxG/YxxD motifs first) would favour interaction with the phagosomal membrane; however, there is also some evidence of EspB being extracted from outer-most layer of the bacterium (Sani *et al.*, 2010). From different experiments (Fig 1), it is expected that oligomerisation is concentration dependent. Interaction with protein structures or a membrane could increase the local concentration, making the system more efficient.

In the third more speculative model, EspB would undergo a conformational change, as observed for some pore-forming proteins such as the amphitropic gasdermins (Liu & Lieberman, 2020). Upon proteolysis, a pre-pore ring could assemble prior to membrane insertion (Ruan *et al.*, 2018). Recently, it was suggested that the PE/PPE family of proteins could form small molecule-selective channels analogous to outer-membrane porins, allowing *M. tuberculosis* to take up nutrients through its almost impermeable cell wall (Wang *et al.*, 2020). Despite evidence, it remains a mystery how such soluble heterodimers would insert into a membrane. We hypothesise that, analogous to the heterodimer EsxA/EsxB where EsxA alone can insert into a membrane in acidic conditions (De Leon *et al.*, 2012), the amphiphilic helices of either PE or PPE alone might insert into the membrane. EspB is fundamentally different from PE/PPE pairs in the sense that its PE and PPE parts are fused into a single protein, joined by one long flexible linker able to adopt multiple conformations (Piton *et al.*, 2020). Unlike the EsxA-EsxB heterodimer, where EsxA would act independently from EsxB upon membrane insertion, the PE moiety of EspB would still be linked to its PPE counterpart even if the latter inserts would itself into a membrane. We speculate that such linker could allow EspB to form tubular-like structures while exchanging PE and PPE domains between different molecules (Fig 7 model 3). Such higher-order oligomers, as described for EspC (Lou *et al.*, 2017) and occasionally also found in our data (Fig EV7), could be a component of the secretion apparatus.

Our hypothesis that EspB acts as a scaffold or structural component of the secretion apparatus is supported by earlier findings. As ESX-1 work through a contact-dependent mechanism and not by secretion of toxins (Conrad *et al.*, 2017), it is possible that the cytotoxic effects on macrophages observed by Chen *et al.* for EspB was the result of an increment in the machinery activity (Chen *et al.*, 2013). Most of the work described here favours model 1 or 2. More evidence needs to be gathered to falsify or verify any of the models. Techniques like *in situ* cryo-electron tomography of infected immune cells could be used to provide visual insight.

In summary, this study reveals factor that prime the oligomerisation of EspB and presents evidence that supports the hypothesis that EspB is a structural element of ESX-1 secretion system, possibly acting on a lipid membrane. ESX-1 is a major player in the virulence of mycobacterial species, like *M. tuberculosis.* However, after decades of arduous research, our understanding on the structure and the mechanism of action of this system remains limited. Here we provide a structural and possibly functional understanding of an ESX-1 element. Full understanding of all the ESX-1 components and structural states could guide structural-based drug and vaccine design in order to tackle the global health threat that tuberculosis is.

## Materials and Methods

### Cloning, expression and purification of EspB constructs

Different constructs used in this study are listed in S1 Table. DNA fragments were PCR-amplified with KOD Hot Start Master Mix (Novagen®) from genomic DNA of *M. tuberculosis* H37Rv, *M. marinum* or *M. smegmatis* [BEI Resources, National Institute of Allergy and Infectious Diseases (NIAID)], and cloned in a modified pRSET backbone (Invitrogen™) using *NsiI* and *HindIII* restriction sites. Constructs included an N-terminal 6×His-tag followed by a tobacco etch virus (TEV) protease cleavage site. EspB mutants and construct EspB_2-385_ and EspB_2-287_ were generated using KOD-Plus-Mutagenesis kit (Toyobo Co., Ltd.) from the plasmid encoding the full-length protein. All plasmids were sequenced to verify absence of inadvertent mutations. *M. haemophilum* and PE25-PPE41 construct were synthesised and codon optimised for expression in *Escherichia coli* (Eurofins Genomics).

For the non-codon optimised constructs, proteins were expressed in Rosetta (DE3) *E. coli* cells in Overnight Express™ Instant LB Medium (EMD Millipore) supplemented with 100 μg/mL of carbenicillin and 25 μg/mL of chloramphenicol for 50 h at 25 °C. In the case of codon optimisation, the protein was expressed in C41 (DE3) *E. coli* cells in the same conditions with the respective antibiotic. Prior to protein purification, cells were resuspended in buffer containing 20 mM Tris-HCl (pH 8.0), 300 mM NaCl, 1 mM PMSF, and 25 U/mL benzonase, and were lysed using an EmulsiFlex-C3 homogenizer (Avestin). Proteins were purified with HisPur™ Ni-NTA Resin (ThermoFisher) equilibrated in the lysis buffer and eluted in the same buffer supplemented with 400 mM imidazole. The 6×His-tag was cleaved using TEV protease followed by a second Ni-NTA purification to remove the free 6xHis-tag, uncleaved protein and the His-tagged protease (Kapust *et al.*, 2001). In case higher purity was needed, proteins were purified on a size-exclusion Superdex200 Increase 10/300 GL column (GE Healthcare) in buffer containing 20 mM Tris-HCl (pH 8.0), 300 mM NaCl. Protein was stored at −80 °C until further use.

### Analytical size exclusion chromatography (SEC)

Samples were dialysed overnight in the corresponding buffers and different concentrations of protein were loaded onto a size-exclusion Superdex200 Increase 3.2/300 column (GE Healthcare Life Science) at a flow rate of 50 μL/min. Basic buffer comprised 20 mM Tris-HCl (pH 8.0), 150 mM NaCl, while the acidic buffer was 20 mM acetate buffer (pH 5.5), 150 mM NaCI.

### Native mass spectrometry

Native mass spectrometry was used to obtain the high resolution mass information of the samples. *M. tuberculosis* EspB_2-348_ (5 mg/mL) was buffer exchanged with 100 mM NH_4_CH_3_CO_2_ (at pH 5.5 and 8.5) using 3-kDa molecular weight cut-off dialysis membrane overnight followed by an extra hour buffer exchange with a fresh NH_4_CH_3_CO_2_ solution at 4 °C. The buffer exchange of fragments produced by limited proteolysis (2 mg/mL) was performed using SEC on a Superdex 200 Increase 3.2/300 column (GE Healthcare Life Science) with 100 mM NH_4_CH_3_CO_2_ at pH 6.8. CH_3_COOH and NH_4_OH were used to adjust the pH of NH_4_CH_3_CO_2_ solution. The mass spectrometry measurements were performed in positive ion mode on an ultra-high mass range (UHMR) Q-Exactive Orbitrap mass spectrometer (Thermo Fisher Scientific) with a static nano-electrospray ionization (nESI) source. Inhouse pulled, gold-coated borosilicate capillaries were used for the sample introduction to the mass spectrometer, and a voltage of 1.2 kV was applied. Mass spectral resolution was set at 4,375 to 8,750 (at m/z=200) and an injection time of 100 to 200 ms was used. For each spectrum, 10 scans were combined, containing 5 to 10 microscans. The inlet capillary temperature was kept at 320 °C. Parameters such as in-source trapping, transfer m/z, detector m/z, trapping gas pressure and mass range were optimized for each analyte separately. All mass spectra were analysed using Thermo Scientific Xcalibur software and spectral deconvolutions were performed with the UniDec software (Marty *et al.*, 2015).

### Cryo-EM sample preparation, data acquisition and image processing

Samples, in 20 mM acetate buffer (pH 5.5), 150 mM NaCl, were diluted to the respective concentrations (Table 1). A volume of 2.5 μL of each sample was applied on glow-discharged UltrAuFoil Au300 R1.2/1.3 grids (Quantifoil), and excess liquid was removed by blotting for 3 s (blot force 5) using filter paper followed by plunge freezing in liquid ethane using a FEI Vitrobot Mark IV at 100% humidity at 4 °C. For PE25-PPE41, an acetate buffer (pH 6.5), 150 mM NaCl was used, due to precipitation of the protein at lower pH.

**Table 1.** Constructs used in this study

| Plasmid name | Species | Gene product | Concentration used for cryo-EM experiments |
| --- | --- | --- | --- |
| pAG01 | *M. tuberculosis* | 6×His-EspB 2-460 | 0.5 mg/mL |
| pAG02 | *M. tuberculosis* | 6×His-EspB 2-358 | 0.5 mg/mL |
| pAG03 | *M. tuberculosis* | 6×His-EspB 2-348 | 0.5 mg/mL |
| pAG04 | *M. tuberculosis* | 6×His-EspB 2-287 | 6 mg/mL |
| pAG05 | *M. tuberculosis* | 6×His-EspB 7-278 | - |
| pAG06 | *M. tuberculosis* | 6×His-MBP-EspB 279-460 | - |
| pAG07 | *M. tuberculosis* | 6×His-EspB 2-348 Q48A | - |
| pAG08 | *M. tuberculosis* | 6×His-EspB 2-348 N55C/T119C | - |
| pAG09 | *M. marinum* | 6×His-EspB 2-355 | 0.5 mg/mL |
| pAG10 | *M. marinum* | 6×His-EspB 2-286 | 8.7 mg/mL |
| pAG11 | *M. haemophilum* | 6×His-EspB 2-287 | 1 mg/mL |
| pAG12 | *M. smegmatis* | 6×His-EspB 2-407 | 10 mg/mL |
| pAG13 | *M. tuberculosis* | PE25 / 6×His-PPE41 | 5 mg/mL |

Cryo-EM single particle analysis (SPA) data were collected using untilted and tilted schemes (Tan *et al.*, 2017). For EspB_2-287_ from *M. tuberculosis* and EspB_2-286_ from *M. marinum*, untilted images were recorded on a Titan Krios at 300 kV with a K3 detector operated in super-resolution counting mode. Tilted SPA data were collected for EspB_2-348_ from *M. tuberculosis* on a 200-kV Tecnai Arctica TEM using SerialEM (Mastronarde, 2005), using a Falcon III detector in counting mode. Table 2 shows all specifications and statistics for the data sets. Individual micrographs of EspB_2-287_ from *M. haemophilum*, EspB_2-348_ from *M. smegmatis* as well as PE25-PPE41 from *M. tuberculosis* were collected on the 200-kV Arctica.

**Table 2.**
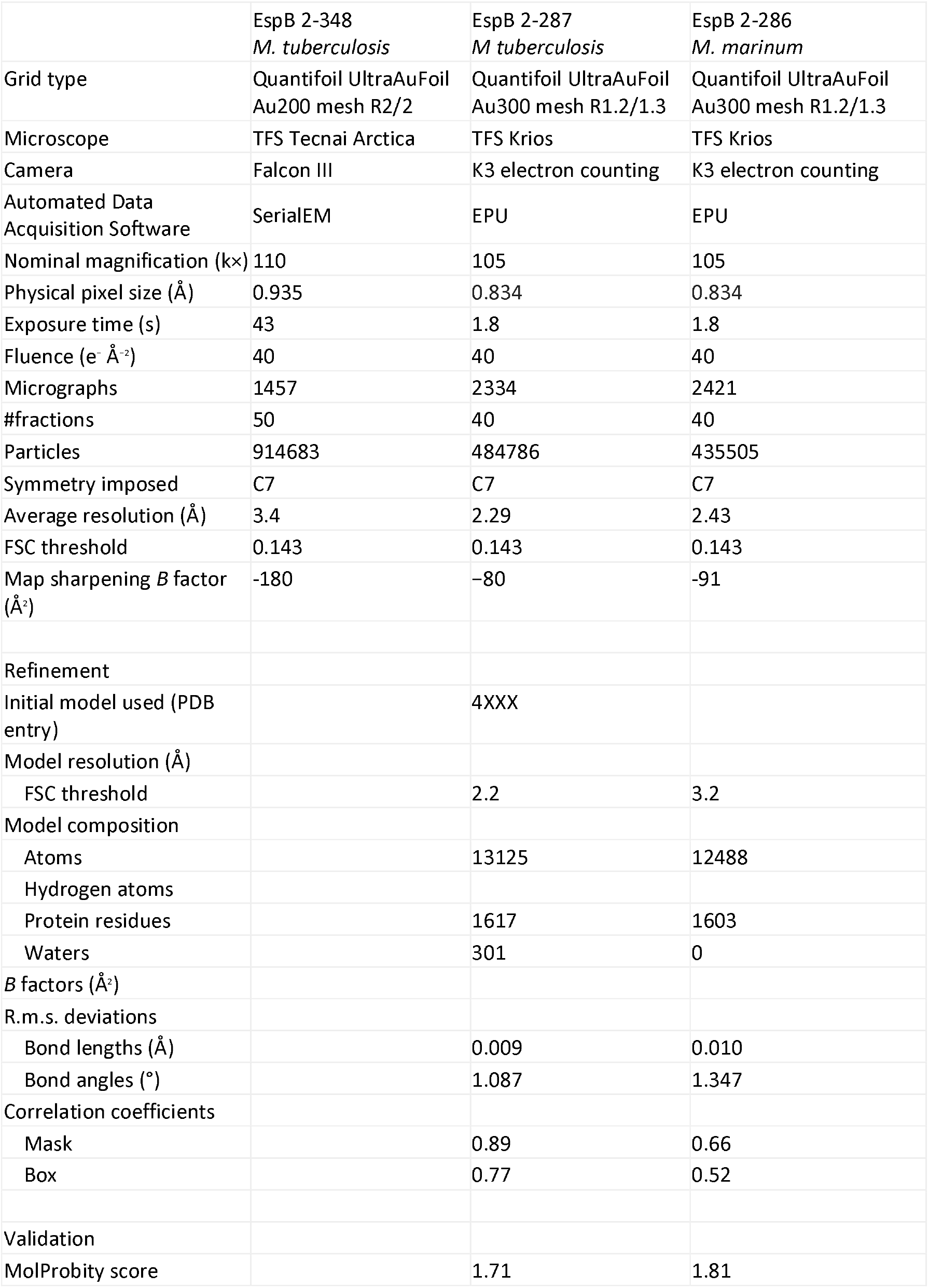
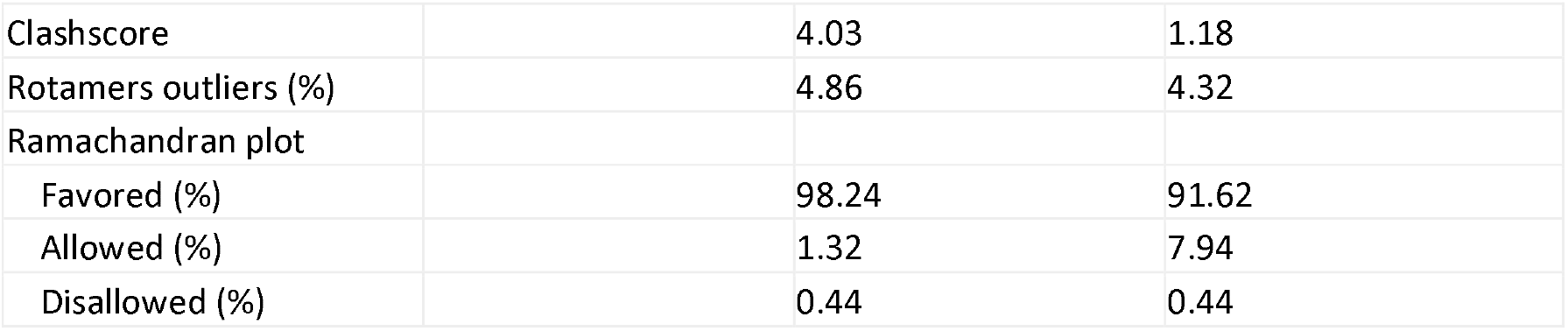
Statistics of cryo-EM data collection, reconstruction and structure refinement

Data were processed using the RELION-3 pipeline (Zivanov *et al.*, 2018). Movie stacks were corrected for drift (5 × 5 patches) and dose-weighted using MotionCor2 (Zheng *et al.*, 2017). The local contrast transfer function (CTF) parameters were determined for the drift-corrected micrographs using Gctf (Zhang, 2016). The EspB_2-348_ data set was collected at two angles of the stage: 0 degrees and 40 degrees. For each tilt angle, a first set of 2D references were generated from manually picked particles in RELION (Scheres, 2012) and these were used for subsequent automatic particle picking. Table 2 lists the number of particles in the final data set after particle picking, 2D classification and 3D classification. The 3D classification was run without imposing symmetry and used to select the heptameric particles. Local CTF parameters were iteratively refined (Zivanov *et al.*, 2018), which was particularly important for the tilted data set, beamtilt parameters were estimated and particles were polished. Particle subtraction followed by focused classification was used to characterise densities other than that described by the refined model described below. Due to extreme preferred orientation of the datasets of EspB_2-348_, automatic masking and automatic B-factor estimation in post-processing were hampered by missing wedge artefacts. For this data set, parameters were manually optimised by visual inspection of the resulting maps. Density within the heptameric pore was obtained by a combination of 2D and 3D classification. The initial density map of a loaded complex was generated by symmetry expansion of a C7 3D-refined particle list, followed by 3D classification in C1 without further image alignment. Later iterations employed 45671 unique particles and 3D refinement in C1 while imposing local symmetry for the heptamer. The resulting 5.3 Å map was used to identify a total of eight EspB monomers (heptamer plus one in the middle), and local symmetry averaged. The final resolution of the heptamer maps, listed in Table 2, varied between 2.3 and 3.4 Å, using the gold-standard FSC=0.143 criterion (Scheres & Chen, 2012).

### Structure determination and refinement

The PDB model 4XXX (Korotkova *et al.*, 2015) was used as a starting model in Coot (Emsley & Cowtan, 2004) for manual docking and building into the tilted-scheme SPA data set of EspB_2-348_ of *M. tuberculosis*. The final model was refined against the high-resolution sharpened map of EspB_2-287_ of *M. tuberculosis*. This model was later used as reference for *M. marinum* model. Models were refined iteratively through rounds of manual adjustment in Coot (Emsley *et al.*, 2010), real space refinement in Phenix (Afonine *et al.*, 2018) and structure validation using MolProbity (Williams *et al.*, 2018).

### Limited proteolysis and Edman sequencing

Samples were incubated with trypsin for different length of time at a molar ratio of 1:6 (enzyme:substrate) following the Proti-Ace™ Kit (Hampton Research) recommendation. The reaction was stopped by adding SDS-PAGE loading buffer (63 mM Tris-HCl, 2% SDS, 10% glycerol, 0.1% bromophenol blue) and samples were resolved on a 12% polyacrylamide gel. Bands were transferred from the SDS-PAGE gel to a PVDF membrane and stained with 0.1% Coomassie Brilliant Blue R-250, 40% methanol, and 10% acetic acid until bands were visible. The membrane was then washed with water and dried, and EspB cleavage products were cut out. The first ten amino acids were determined by Edman sequencing at the Plateforme Protéomique PISSARO IRIB at the Université de Rouen.

### Circular dichroism spectroscopy

The CD spectra of 5 μM EspB_279-460_ were recorded either in 50 mM phosphate (pH 8.0), 50 mM NaCl or 10 mM acetate (pH 5.5), 50 mM NaCl at 25 °C in the far-UV region using a Jasco J-1500 CD spectropolarimeter (JASCO Analytical Instruments) on a 0.1 cm path-length cell. Spectra correspond to the average of five repetitive scans acquired every 1 nm with 5-s average time per point and 1-nm band pass. Temperature was regulated with a Peltier temperature-controlled cell holder. Data were corrected by subtracting the CD signal of the buffer over the same wavelength region. The effect of 2,2,2-trifluoroethanol (TFE) was recorded using the aforementioned phosphate buffer. Secondary structure content was estimated by deconvolution using the program BeStSel (Micsonai *et al.*, 2018).

## Supporting information

Expanded View Figure 1

Expanded View Figure 2

Expanded View Figure 3

Expanded View Figure 4

Expanded View Figure 5

Expanded View Figure 6

## Data availability

The final maps as well as the half-maps and masks will be deposited in EMPIAR. The refined *M. tuberculosis* and *M. marinum* will be deposited within the Protein Data Bank.

## Acknowledgments

We thank Paul van Schayck (UM) for indispensable SerialEM and IT support; the Microscopy CORE Lab (UM) for their technical and scientific support; Yue Zhang (UM) for help in model refinement; Chris Lewis (UM) for help with the tomograms; Laurent Coquet (Université de Rouen, France) for Edman sequencing; Florence Pojer and Stewart Cole (Global Health Institute, Lausanne, Switzerland) for initial sample aliquots and preliminary studies; Ron Heeren and Shane Ellis (UM) for native mass spectrometry support; and Hang Nguyen (UM) for critical reading of the manuscript. This research received funding from the Netherlands Organisation for Scientific Research (NWO) in the framework of the Fund New Chemical Innovations, numbers 731.016.407 and 184.034.014, from the European Union’s Horizon 2020 Research and Innovation Programme under Grant Agreement No 766970 Q-SORT. This research is also part of the M4I research programme supported by the Dutch Province of Limburg through the LINK programme.

## Author Contributions

AG and RBGR designed the study and wrote the manuscript. AG, VV, YG, GT, AS, NSP and AM performed the experiments. AG, GT, NSP and RBGR analysed the data. CLP, PJP and RBGR supervised the project. All authors read and approved the final manuscript.

## Expanded View Figure legends

**Fig EV1. Sequence alignment of EspB from different mycobacterial species.** Numbering and sequence identity are based on the sequence of *M. tuberculosis*. Alignment was generated using ClustalW server, and figure was created using Jalview software (Waterhouse *et al.*, 2009). The colour scheme of ClustalX is used (Larkin *et al.*, 2007).

**Fig EV2. EspB preferential orientation caused by an interaction to the air-water interface.** (A-B) Tomogram slice of EspB_2-348_ with 27 nm thickness in X,Y and X,Z orientation, respectively.

**Fig EV3. Cryo-EM analysis of EspB structure.** Gold-standard Fourier shell correlation (FSC) plot of EspB_2-287_ from *M. tuberculosis* (A) and EspB_2-286_ from *M. marinum* (B). (C) Quality of cryo-EM-derived density map. Selected regions showing the fit of the derived atomic model to the cryo-EM density map (black mesh) (D) Size exclusion chromatograms of EspB_2-348_ from *M. tuberculosis* and mutants that affect oligomerisation.

**Fig EV4. Oligomerisation of EspB is independent of the integrity of the PE-PPE linker.** (A) Trypsin digestion of different EspB constructs from *M. tuberculosis* over 1-4 h. (B) Structural model of an EspB monomer (PDB ID 4XXX) showing the PE-region (gold), PPE-region (grey) and the trypsin cleavage site (arrow, residues R121-V112). (C, D) Native mass spectrometry of the trypsin-digested sample, raw and deconvoluted data. Colour coding as in panel (B). Inset table compares the respective mass of the fragments calculated from the sequence and native mass spectrometry. (E) SEC (top) and SDS-PAGE (bottom) of undigested (black) and trypsin-digested (red) EspB_2-348_.

**Fig EV5. Characterisation of the C-terminal region of EspB.** (A) Kyte-Doolittle hydrophobicity plot of residues 280-460 of EspB from *M. tuberculosis.* Inset shows the degree of hydrophobicity of residues 280-360 from different species. Window size of 9 was used as parameter. (B, C) Far UV circular dichroism spectra of *M. tuberculosis* EspB_279-460_ at different pH and TFE concentrations. Inset in (B) shows the spectrum-difference between pH 5.5 and pH 8.0.

**Fig EV6. Higher-order oligomer formation.** Size exclusion chromatography profiles of EspB_2-348_ from *M. tuberculosis* (20 mg/mL) injected onto a Superdex200 Increase 10/300 GL. Inset corresponds to a Blue-Native PAGE of the SEC fractions highlighted in red.

