## Supplementary figures and images for "Priming mycobacterial ESX-secreted protein B to form a channel-like structure"

### Expanded View Figure 1

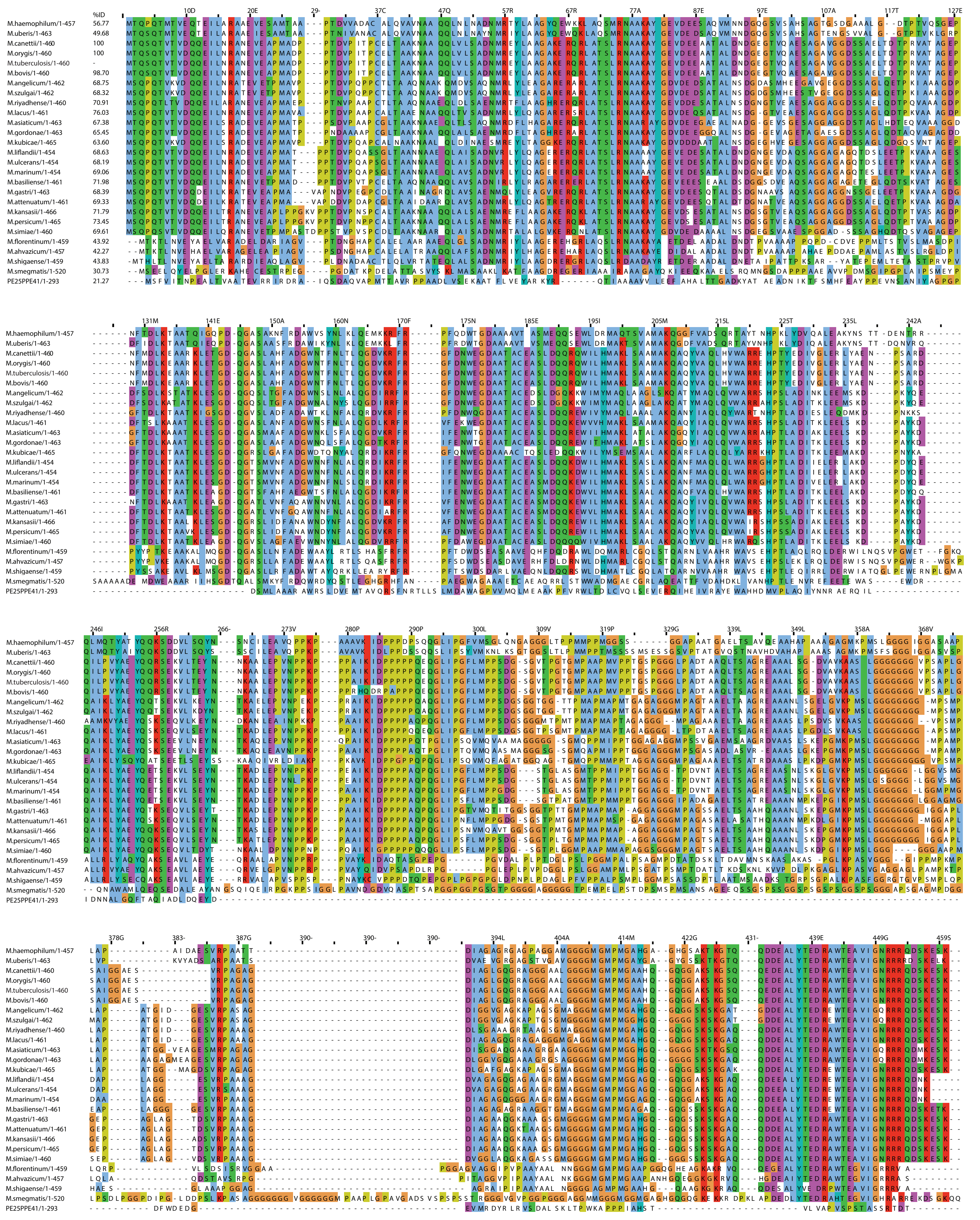

### Expanded View Figure 2

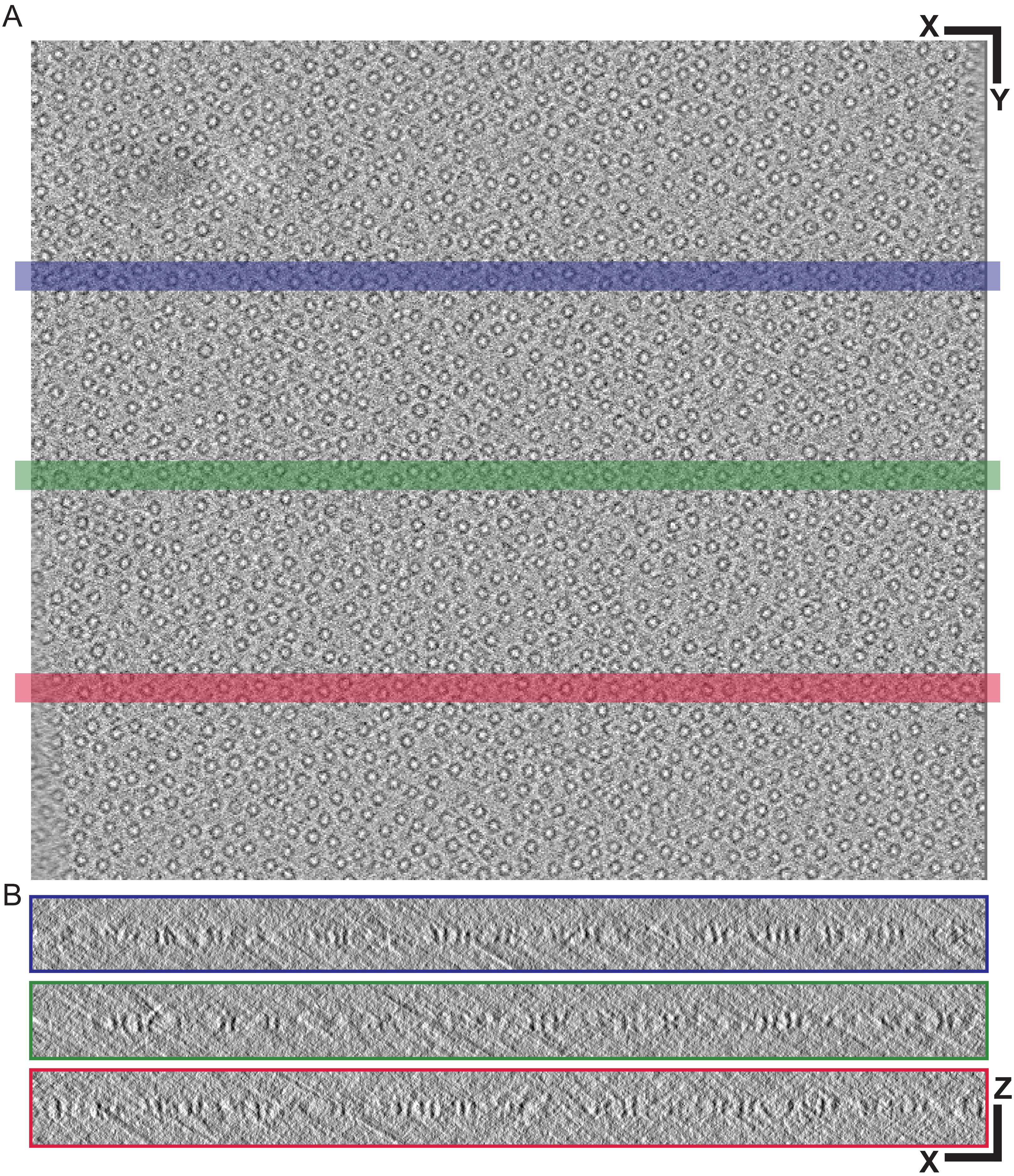

### Expanded View Figure 3

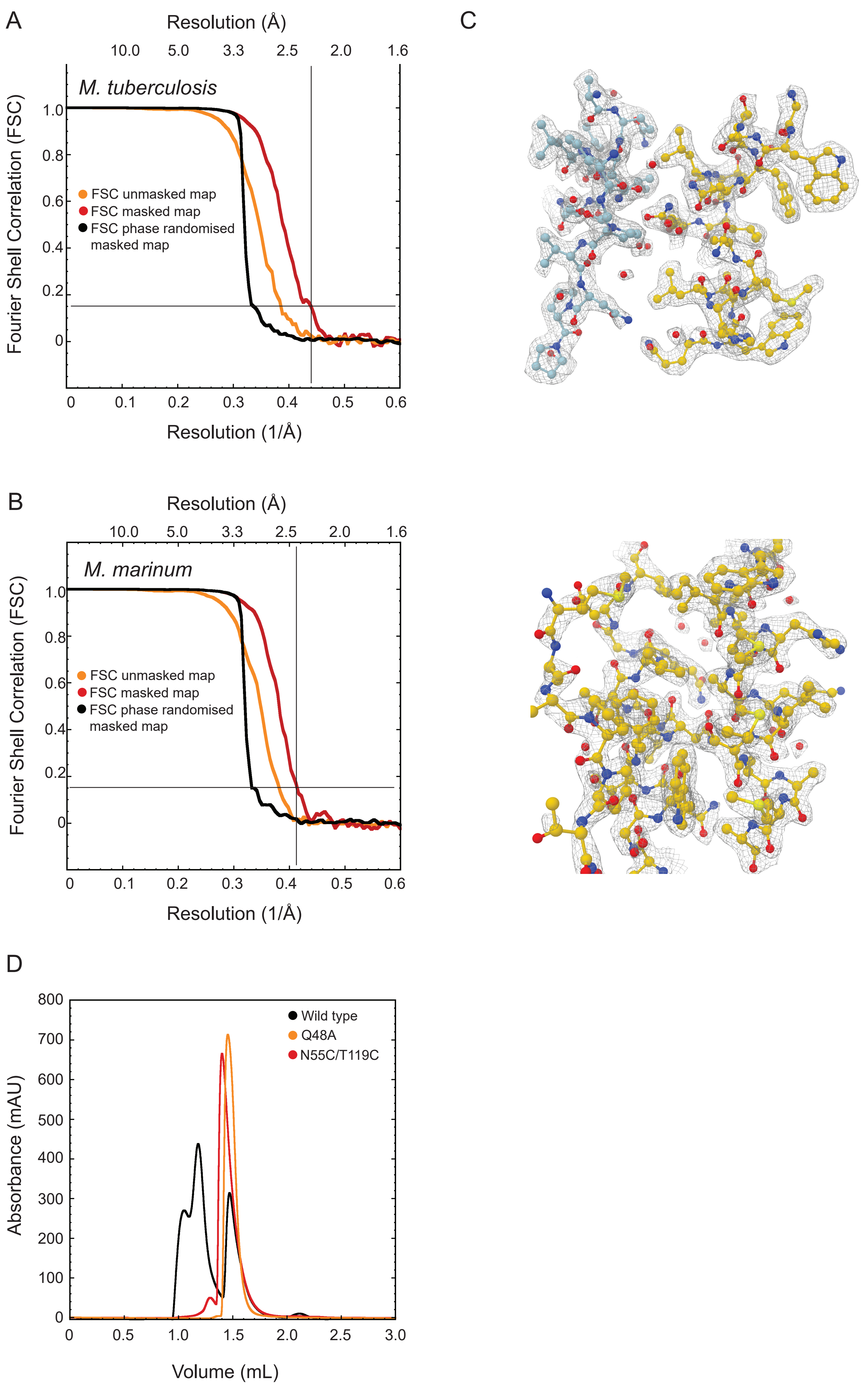

### Expanded View Figure 4

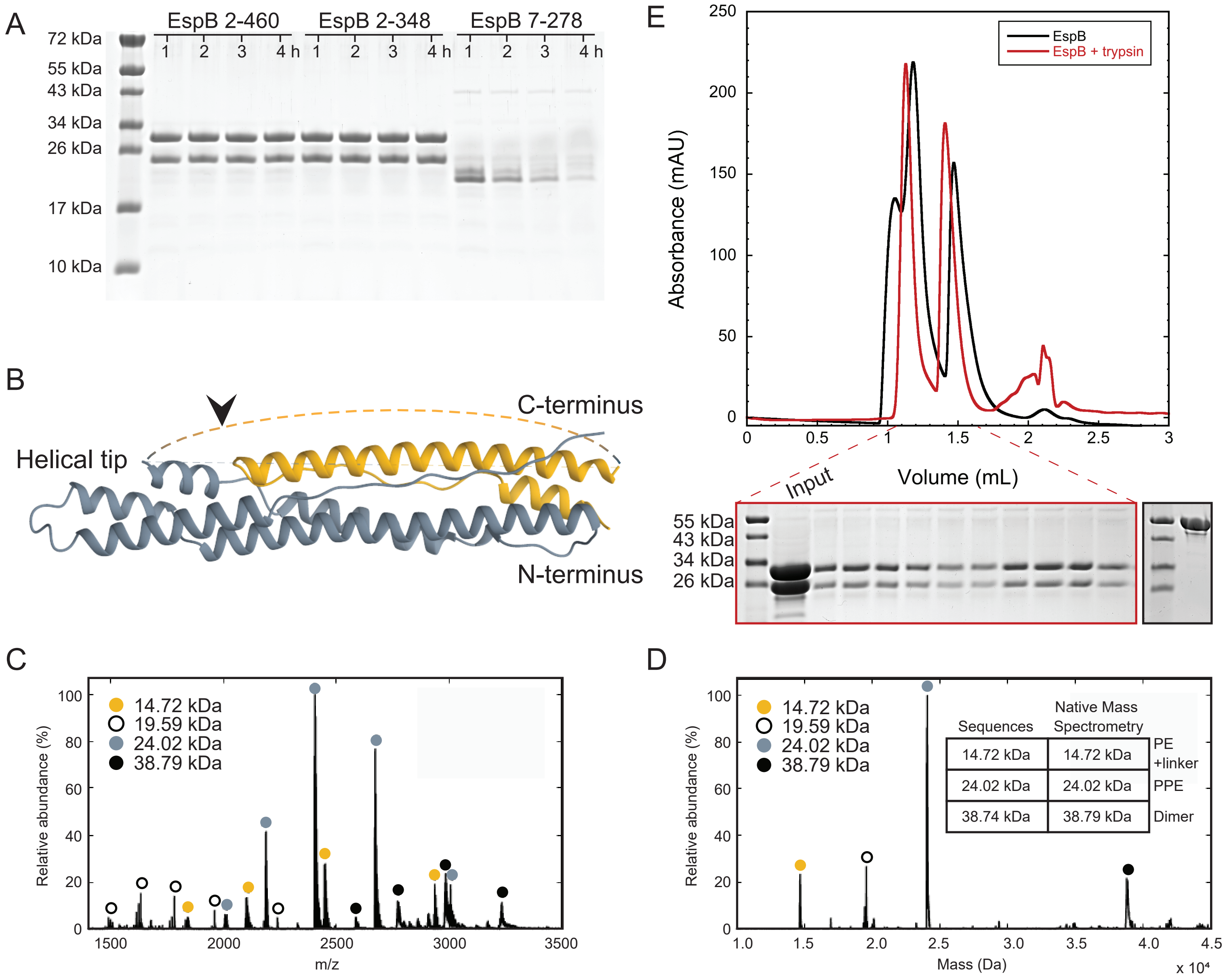

### Expanded View Figure 5

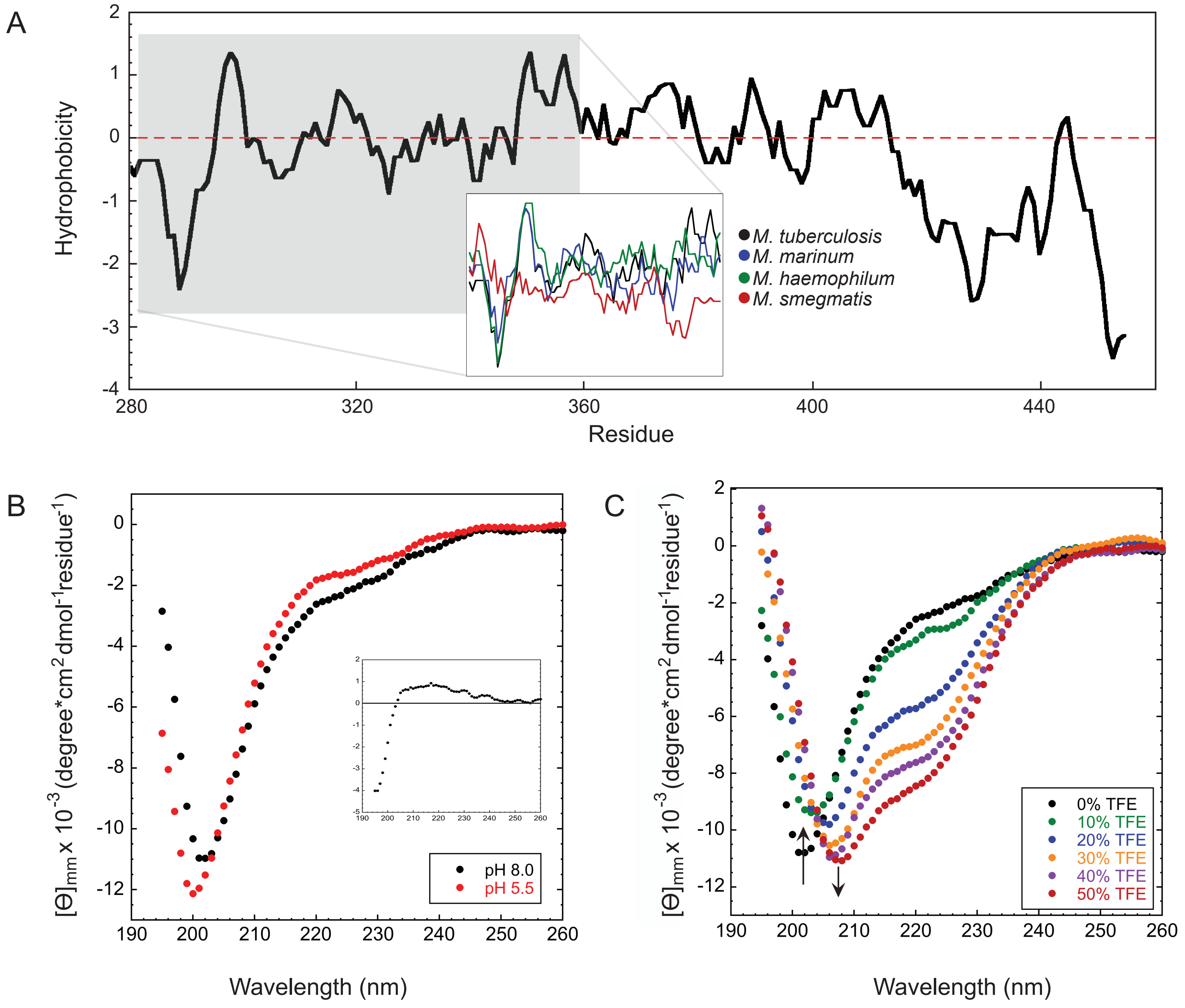

### Expanded View Figure 6

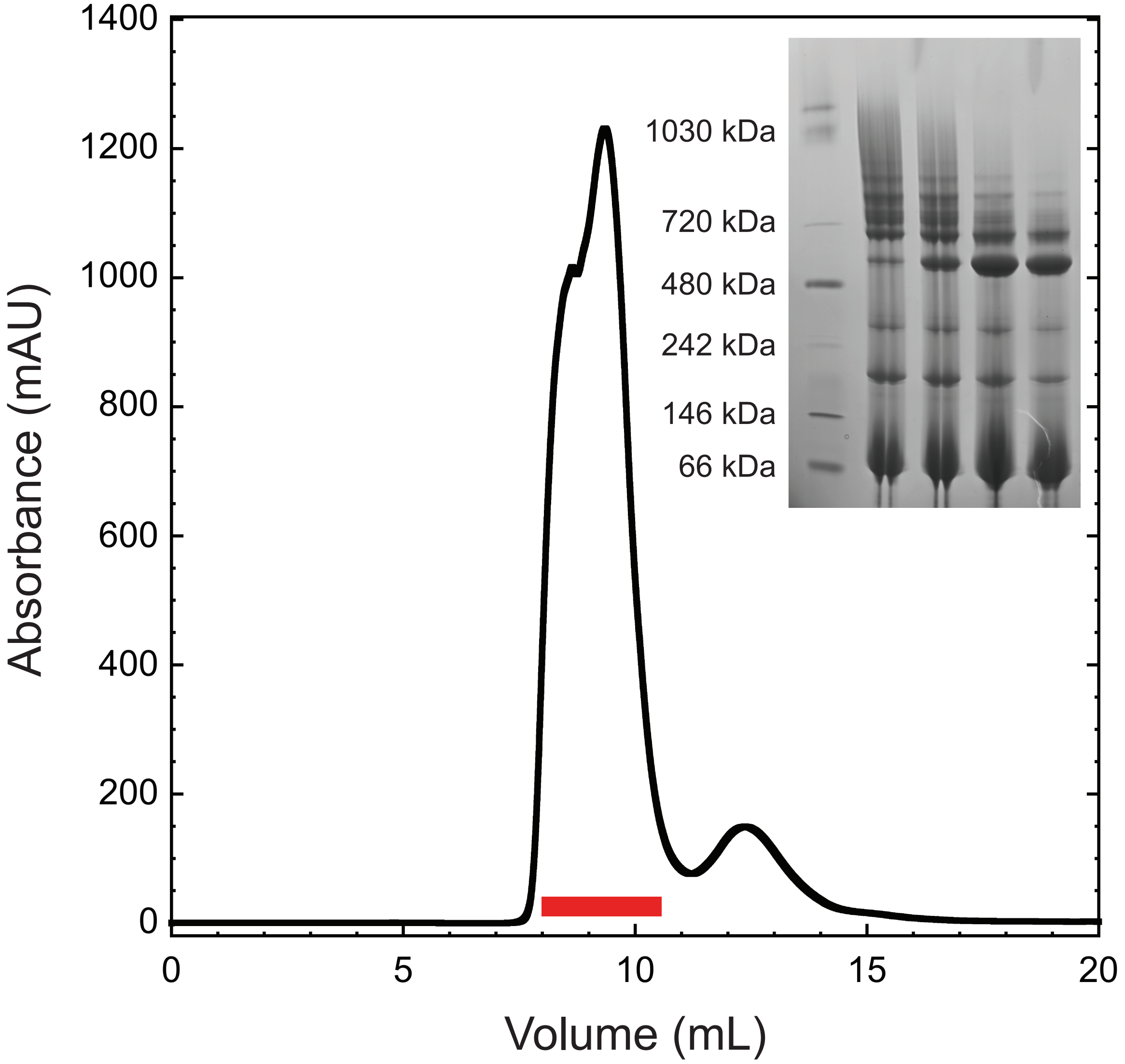
